# Sequential single cell transcriptional and protein marker profiling reveals TIGIT as a marker of CD19 CAR-T cell dysfunction in patients with non-Hodgkin’s lymphoma

**DOI:** 10.1101/2021.04.26.441326

**Authors:** Zachary Jackson, Changjin Hong, Robert Schauner, Boro Dropulic, Paolo F. Caimi, Marcos de Lima, Kalpana Gupta, Jane S. Reese, Tae Hyun Hwang, David N. Wald

## Abstract

Chimeric antigen receptor T cell (CAR-T cell) therapy is known to produce durable remissions in the treatment of CD19^+^ relapsed/refractory B cell malignancies. Nonetheless, a significant portion of patients receiving the therapy experience poor outcomes in the acute response for unknown reasons. Given the decreased expansion and persistence of CD8 CAR-T cells in poor outcome groups, this failure may be attributed to CAR-T cell dysfunction. However, a comparison of the post-infusion transcriptional profiles and phenotypes between CAR-T cells of poor and favorable response groups has not been performed. Here, we employed single cell RNA sequencing and protein surface marker profiling of serial CAR-T cell blood samples from patients with CD19^+^ relapsed/refractory non-Hodgkin’s lymphoma (NHL) to reveal CAR-T cell evolution, identify biomarkers of response, and test for evidence of exhaustion in CAR-T cells of poor responders. At the transcriptional and protein levels, we note the evolution of a majority of CAR-T cells toward a non-proliferative and highly-differentiated state. In poor outcome patients, we observed a more marked enrichment of an exhaustion profile as compared to favorable outcome patients. Lastly, we identified the checkpoint receptor TIGIT (T cell immunoreceptor with Ig and ITIM domains) as a novel prognostic biomarker and potential driver of CAR-T cell exhaustion. Altogether, we provide evidence of CAR-T cell dysfunction marked by TIGIT expression driving poor response in NHL patients.

## Introduction

CAR-T cells are T cells engineered with a chimeric antigen receptor to specifically lyse tumor cells expressing the targeted antigen. The safety and efficacy of CD19 CAR T cell products in B cell malignancies has led to FDA authorization of four products for treatment of pediatric B cell acute lymphoblastic leukemia (B-ALL) and subtypes of non-Hodgkin’s lymphoma (NHL). The CAR construct designs used in these applications contain intracellular CD3ζ with either 4-1BB (4-1BB.CAR) or CD28 (CD28.CAR). The 4-1BB intracellular domain is thought to convey resistance to exhaustion and superior CAR-T cell persistence in comparison to the CD28 intracellular domain which exhibits greater short term activity and adverse events^1–3^. Here, a 4-1BB design was applied in the generation of CAR-T cells for the treatment of relapsed/refractory non-Hodgkin’s lymphoma (NHL).

While approximately 50% of patients with NHL undergo remission after initial chemotherapy, relapsed/refractory disease is essentially incurable without intervention with adoptive cell therapies^4–7^. Following CD19 CAR-T cell therapy, approximately 30-40% of NHL patients will have durable remissions^8^, ^9^. Mechanisms of early relapse or refractory disease remain inconclusive, with significant variation between studies due to differences such as disease type, product design, and product manufacture^10–14^. Proposed mechanisms of resistance include poor T cell quality, T cell exhaustion, antigen loss/modulation, host factors, and tumor microenvironment^15^. While several studies have focused on comparisons of infusion products, detailed single cell studies evaluating post-infusion CAR-T cell transcriptional profiles and phenotypes associated with clinical outcomes remain lacking^16, 17^. We address this gap by evaluating CAR-T cells from serial pre- and post-infusion samples in patients with both favorable and poor outcomes by applying an innovative method combining single cell transcriptomics and cell surface protein expression in individual cells.

## Results

### CD19 CAR-T cells demonstrate significant transcriptional heterogeneity that changes after infusion into patients

To describe the evolution of CAR-T cells after infusion into NHL patients and to identify mechanisms and biomarkers of response, our study examined manufactured CAR-T cell products and isolated CAR-T cells from post-infusion blood samples from patients treated for CD19^+^ relapsed/refractory NHL (Supplemental Table 1). Utilizing scRNA sequencing and flow cytometry, we investigated time points after infusion that are known from previous studies to be associated with peak expansion (day 14) and contraction (day 30) and represent key changes in CAR-T cell activity^18^ (study schema depicted in Figure 1A). Altogether, our datasets include 14 manufactured CAR-T cell products, 13 samples from day 14, and 12 samples from day 30. This sampling represents 10 patients with favorable response (complete or partial remission (CR; PR)) and 4 patients with poor response (stable or progressive disease (SD; PD)) (Supplemental Table 2).

**Figure 1.**
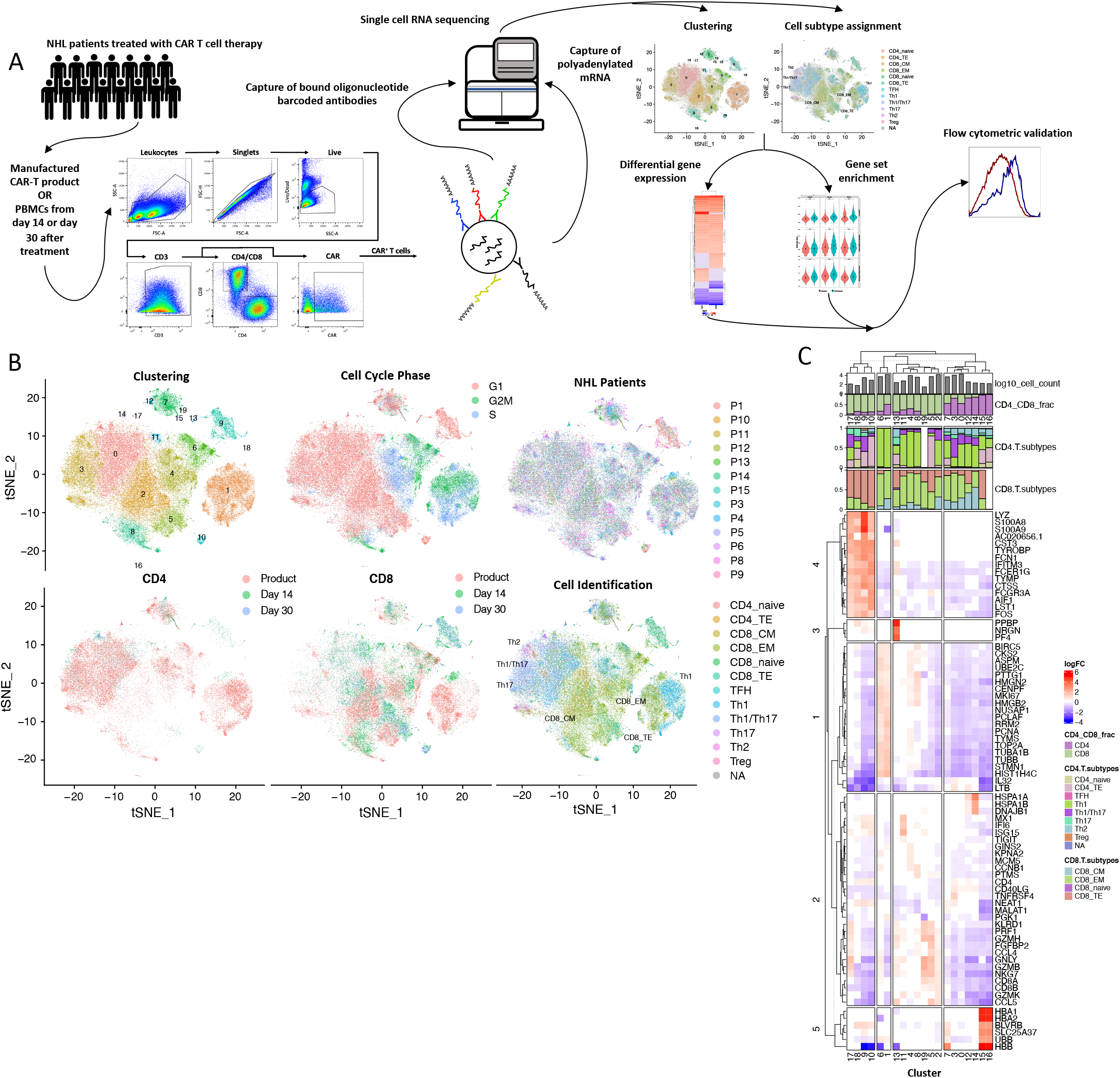
CD19 CAR-T cells demonstrate significant transcriptional heterogeneity that changes after infusion into patients. **A)** Study scheme. Viable CAR-T cells were sorted from the CAR-T cell products or PBMCs of NHL patients. Single cell libraries for captured CAR-T cell mRNA and feature barcoding transcripts were then prepared with the 10x Genomics Chromium single cell 3’ platform with feature barcoding technology and subsequently sequenced. After quality control, dimension reduction was performed and analyzed by cluster or cell subtype with differential gene expression and gene set signature scoring. Validation of memory marker and exhaustion marker expression was performed with flow cytometry. **B)** tSNE based on scRNA overlays depicting cluster assignment, cell cycle analysis, patient number, CD4 or CD8 T cell group by time point, and T-cell subtype. **C)** Heatmap of differentially expressed genes (adjusted p-value < 0.05) between all clusters of B) by log fold change. Only top 3 to 10 genes with highest absolute log fold change are represented. Clusters are annotated with cell subtype proportions.

To isolate CAR-T cells for scRNA sequencing, viable CD3^+^CAR^+^ cells were sorted from cryopreserved CAR-T cell products or PBMCs. Next, libraries were generated with the 10x Genomics Chromium single cell 3’ platform with feature barcoding technology to allow simultaneous and paired quantification of transcriptional and cell surface protein expression in individual CAR-T cells^19^. The inclusion of feature barcoding in addition to enabling assessments of key markers at the protein level also allowed the discrimination of the memory markers CD45RA and CD45RO that cannot be discriminated at the RNA level. The libraries were sequenced and the data stringently filtered to remove on average 8.1% mitochondrial reads and yielded 94,000 cells with an average of 3,917 cells per sample, 8,518 reads per cell, and 2263 unique detectable genes per cell (Supplemental Table 3). Batch effect removal was applied to remove differences due to sample preparation or sequencing.

To appreciate the heterogeneity present in the CAR-T cells both among patients and time points, RNA-based dimension reduction and unbiased clustering was applied on the scRNAseq dataset to yield 19 clusters with distinct transcriptional profiles (Figure 1B;C). At pre-infusion, a consistent pattern of clustering was observed across patients, with the predominant cluster similar in 9 out of 12 patients (C1). After infusion, heterogeneity increased across patients, with different predominant clusters in 4 of 7 patients at day 14 and 4 of 5 patients at day 30 (Supplemental Figure 1A). In accordance with peak expansion expected around day 14, clusters primarily composed of pre-infusion and day 14 samples (C1, C4, and C6) contained the most actively-dividing cells as evidenced by their abundance in the S phase by cell cycle analysis (Figure 1B; Supplemental Figure 1B). Differential gene expression of individual clusters compared to other clusters demonstrated the cells belonging to these clusters were highly enriched (FDR < 0.05) in genes associated with immature T cell types and proliferation (Figure 1C)^20^. On the other hand, other major clusters (C2, C5, C8) which consist predominantly of CD8 CAR-T cells from day 14 and day 30 samples were found not to be actively proliferating and exhibited higher expression (FDR < 0.05) of genes associated with effector CD8 T cell phenotypes including GZMB, GZMH, GZMK, PRF1, GNLY, CCL4, and CCL5 (Figure 1C). To investigate if the heterogeneity across clusters correlated with particular T cell subtypes, SingleR cell ID annotation was applied. SingleR utilizes predefined T cell subtype gene signatures to assign a CD4 or CD8 T cell subtype to each cell, including naive, central memory (CM), effector memory (EM) and terminal effector (TE) CD8 T cell subtypes^21, 22^. Application of cell identification results to the dimension reduction plot showed distinct localization of CD4 and CD8 T cells. While the predominant CD8 subtype was effector memory, two clusters were enriched with central memory transcriptional profiles (C0, C2) and three clusters enriched with terminal effector transcriptional profiles (C5, C9, C10) (Supplemental Figure 1C). Overall, CART cells demonstrated significant heterogeneity across time points (product, day 14, day 30), cell cycle phase, cell type, and patient.

### Circulating CD8 CAR-T cells differentiate to an effector-like state and express high levels of TIGIT post-infusion

Detailed descriptions of changes in CAR-T cell transcriptional profiles and surface marker expression patterns after infusion remain lacking. Consistent with previous reports that clinical responses are driven by the expansion of cytotoxic CD8 T cells, we observed that the relative proportion of CD8 to CD4 CAR-T cells significantly increased after infusion from 52% to 87% CD8 on average (p < 0.001) (Figure 2A)^23^. Given this observation and the greater representation of CD8 CAR-T cells in our samples, our analyses focused on CD8 CAR-T cells. The most significantly upregulated genes in post-infusion CD8 CAR-T cells included transcription factors (PRDM1, EOMES) and cytotoxic effector molecules (GZMB, PRF1, GZMK, CCL5) associated with differentiation into cytotoxic effector cells (padj < 0.05) (Figure 2B). Notably, transcription factors associated with exhaustion (TOX, TOX2, NR4A2, NR4A3) were also significantly upregulated post-infusion, and gene set enrichment analysis of memory/effector versus exhaustion gene sets showed post-infusion CD8 CAR-T cells were significantly enriched in exhaustion-related genes compared to pre-infusion (p < 0.05) (Supplemental Figure 2A). Differential gene expression of pre- and post-infusion CD8 CAR-T cells within each of the clusters demonstrated these changes were largely independent of clustering (Figure 2C). Notably, upregulated expression of all exhaustion marker genes investigated in post-infusion CD8 CAR-T cells was observed, with TIGIT surprisingly being the most significant (pre- vs. post-infusion; log2FC = 2.39; p < 0.0001) (Figure 2D). Quantification of cell type proportions across time points using SingleR transcription-based cell assignments indicated a shift from an equal proportion of CD8 central memory and effector memory profiles in the product towards an effector memory profile at day 14 and a combination of effector memory and terminal effector profiles at day 30 (Figure 2E). Altogether, gene expression analyses indicate CD8 CAR-T cells undergo differentiation towards a cytotoxic effector profile and exhaustion after infusion into patients.

**Figure 2.**
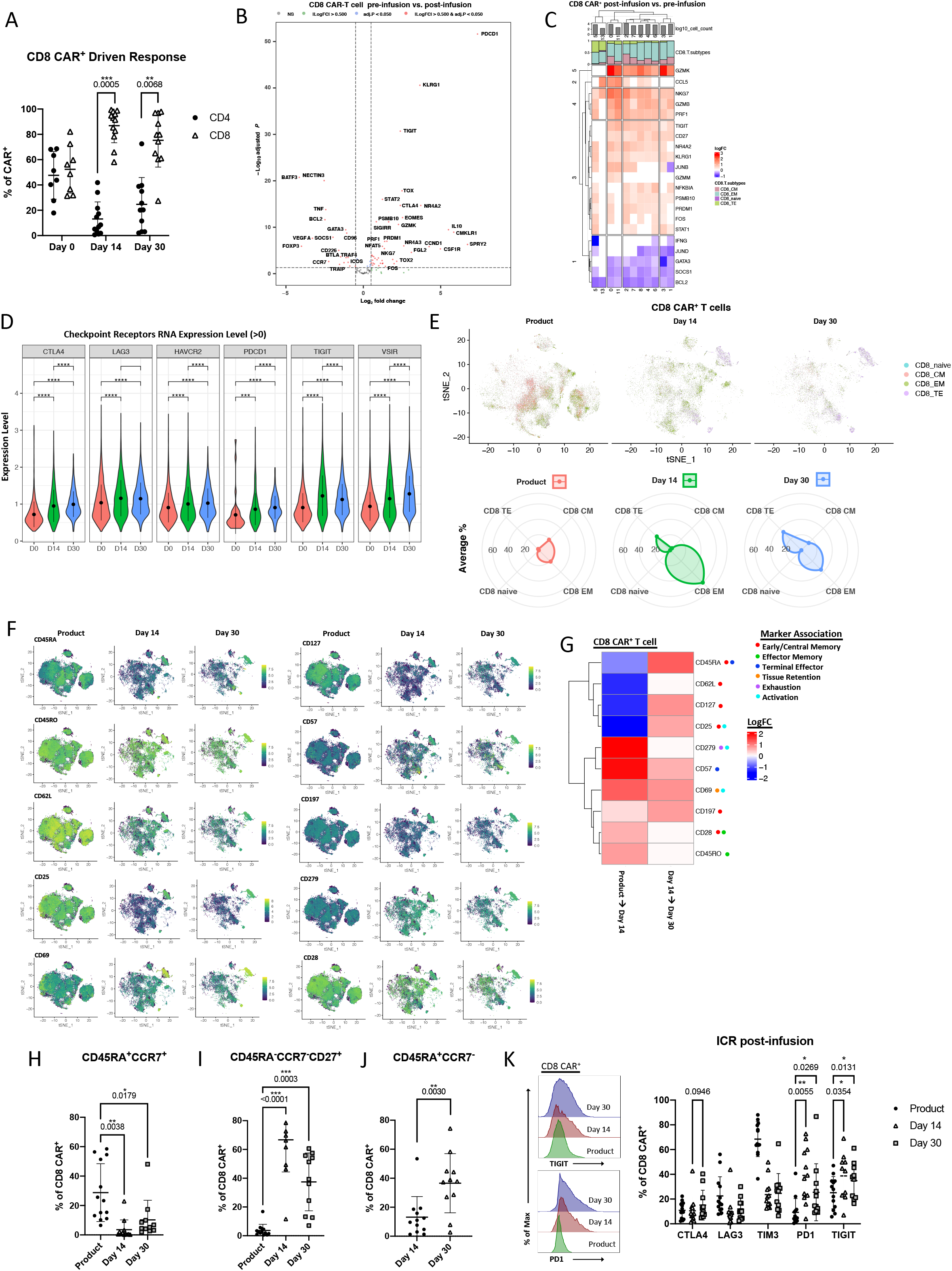
Circulating CD8 CAR-T cells differentiate to an effector-like state and express high levels of TIGIT post-infusion. **A)** Percentage of CD4 or CD8 CAR-T cells of total CAR-T cells across time points by flow cytometry. **B)** Volcano plot of differentially expressed genes (adjusted p-value < 0.05) in post-infusion CD8 CAR-T cells compared to product CD8 CAR-T cells. **C)** Heatmap of differentially expressed genes (adjusted p-value < 0.05) from an immunoregulatory gene list between pre- and post-infusion CD8 CAR-T cells within individual clusters by scRNA seq. Color represents average log fold change in post-infusion cells. **D)** Log normalized gene expressions of exhaustion markers are compared across time points by Wilcoxon rank-sum test. Only cells with the expressed gene are considered. **E)** Top – scRNA tSNE dimension reduction of all samples with CD8 cell subtype assignment overlay across time points. CD4 CAR-T cells excluded from plot. Bottom – relative frequency of CD8 T cell subtype assignments of total CAR-T cells at each time point. **F)** scRNA tSNE dimension reduction plots with relative surface expression overlay for each of the indicated markers as measured by feature barcoding. Each of the indicated time points contains total CAR-T cells from that time point. **G)** Heatmap of differentially expressed surface markers (adjusted p-value < 0.05) from F) in CD8 CAR-T cells between time points. Color represents log fold increase/decrease in the latter time point. **H-J)** Percentage of CD8 CAR-T cells with the indicated memory phenotype across time points as measured by flow cytometry. **K)** Left – Histograms of fluorescence intensity of TIGIT and PD1 across time points as measured by flow cytometry. Each curve represents a concatenation of all samples with equal proportions of CD8 CAR-T cells from each sample. Right – Comparison across time points of the percentage of CD8 CAR-T cells expressing checkpoint receptors CTLA4, LAG3, TIM3, PD1, or TIGIT as measured by flow cytometry.

Given the changes in the transcriptional profile of CD8 CAR-T cells post-infusion, changes in cell surface phenotype upon infusion were expected to reflect differentiation towards effector memory or terminally differentiated phenotypes. We have previously demonstrated that CD8 CAR-T cell products are enriched in an early memory phenotype with expression of CD45RA, CCR7, and TCF7^24^. Here, feature barcoding markers were chosen for their association with stemness/early memory (CD45RA, CD197 (CCR7), CD127 (IL-7R), CD62L, CD25, CD28), activation/effector memory (CD25, CD69, CD45RO, CD279 (PD1)), terminal differentiation (CD45RA, CD57) and exhaustion (PD1)^25, 26^. Consistent with the RNA expression data, we observed greater surface expression of the naive and early memory markers CD45RA, CD127, CD62L, and CD25 in the product (Figure 2F;G). By day 14, a global increase in CD45RO, CD69, CD57, and PD1 (CD279) was observed suggesting differentiation toward an activated, effector-like state (Figure 2F;G). However, at the cluster-level, clusters 5, 9, and 10 were found to be enriched in CD45RA and CD57, indicative of cluster-specific enrichment with a terminally differentiated phenotype. We next compared changes from day 14 to day 30 for evidence of further differentiation and observed an additional increase in global CD45RA and CD57 expression. On the other hand, there was a global increase in the memory markers CD127, CD25, and CD197 from day 14 to day 30, consistent with contraction of effector cells by day 30 after tumor resolution. Overall, the cell surface phenotypes of CD8 CAR-T cells both pre- and post-infusion as measured by feature barcoding showed high similarity to the RNA expression results indicating CD8 T cell differentiation to effector memory and terminal effector phenotypes.

Next flow cytometry was utilized to validate changes in memory status and exhaustion marker expression in CD8 CAR-T cells with additional patient samples (Supplemental Table 2). Similar to the previous analysis using surface protein feature barcoding, we observed an increased proportion of CD45RA^+^CCR7^+^ cells in the product (average 29%) as compared to post-infusion samples (average 3.6%), indicative of a greater proportion of early memory T cells (Figure 2H). At day 14, an average 83.3% of CD8 CAR-T cells were CD45RA^−^ and the predominant phenotype (average 66.7%) was CD45RA^−^CCR7^−^CD27^+^, a significant shift toward an effector phenotype as compared to CAR-T cells in the product (p < 0.0001) (Figure 2I). While day 30 samples also contained high frequencies of CD45RA^−^CCR7^−^CD27^+^ cells, there was a significant increase in CD45RA^+^CCR7^−^ cells among CD8 CAR-T cells, once again indicative of terminal differentiation (p< 0.01) (Figures 2J;Supplemental Figure 2B). Notably, at both day 14 and day 30, CD8 CAR-T cells could be distinguished from endogenous CD8 T cells in our flow cytometry dataset by expression of CD27, which may contribute to the long-term maintenance observed in 4-1BB.CAR-T cells (Supplemental Figure 2C)^27^. Altogether, the data supports a shift from early memory to effector memory and terminal effector CD8 CAR-T cells in post-infusion samples.

As gene expression and feature barcoding datasets indicated higher levels of exhaustion marker expression post-infusion, we further assessed for changes in checkpoint molecules by flow cytometry. In the manufactured CAR-T cell product, 68% of CD8 CAR-T cells expressed TIM3. While this could be indicative of early exhaustion or senescence, this expression was attributed to activation during CAR-T cell manufacture, as expression dropped after infusion to an average 24% TIM3^+^ (Figure 2K). Among the exhaustion markers assayed, PD1 and TIGIT expression were significantly induced, and TIGIT expression was the most sustained at day 30 (p < 0.05) (Figure 2K). To determine if the observed expression was attributable to direct CD19 antigen encounter, we compared expression of exhaustion markers in CD8 CAR-T cells to endogenous CD8 T cells. CD8 CAR-T cells showed enrichment of exhaustion markers, and the most prominent were CTLA4, TIM3, and TIGIT (Supplemental Figure 2D). Altogether, our data indicates elevated expression of TIGIT in CD8 CAR-T cells post-infusion, and TIGIT expression in these cells is likely at least partially due to direct antigen encounter.

### CAR-T cells of poor responders are enriched in an exhaustion-like phenotype post-infusion with high TIGIT expression

With evidence of exhaustion in post-infusion CD8 CAR-T cells, we next investigated for differences in the exhaustion profile in CAR-T cells from favorable and poor responders. We found CD8 CAR-T cells of poor responders exhibited significantly decreased expansion and persistence compared to responders by flow cytometry (p < 0.05) (Figure 3A). At the RNA cluster-level, cell frequencies across responder groups were comparable (Supplemental Figure 1A, Figure 3B). We next compared differentially expressed genes of post-infusion CD8 CAR-T cells between poor and favorable response groups to identify dysregulated genes in the CD8 CAR-T cells of poor responders. Of note, significantly upregulated transcription factors included FOS, JUNB, JUND, FOSB, JUN, NR4A2, NFKBIA, and PRDM1(FDR < 0.0001) (Figure 3C). These changes were also largely consistent across clusters (Figure 3D). To confirm a dysfunctional profile in CAR^+^ CD8 T cells from poor response patients, we performed dysfunction scoring by applying three different exhaustion signatures^1, 28, 29^. Applying these signatures to post-infusion CD8 CAR-T cells we consistently observed significantly higher dysfunctional scores in CD8 CAR-T cells of poor responders both globally (p < 0.0001) and within the most predominant clusters (Figure 3E). Higher dysfunction scores were also consistent across CD8 CAR-T cell types (Supplemental Figure 3A). Upon direct investigation of exhaustion marker expression within individual patients, a greater upregulation of TIGIT was observed in the CD8 CAR-T cells of poor responders (logFC = 0.7) as compared to responders (logFC = 0.5) (Figure 3F). This was also evident in the percentage of CD8 CAR-T cells expressing TIGIT at the RNA level (Figure 3G).

**Figure 3.**
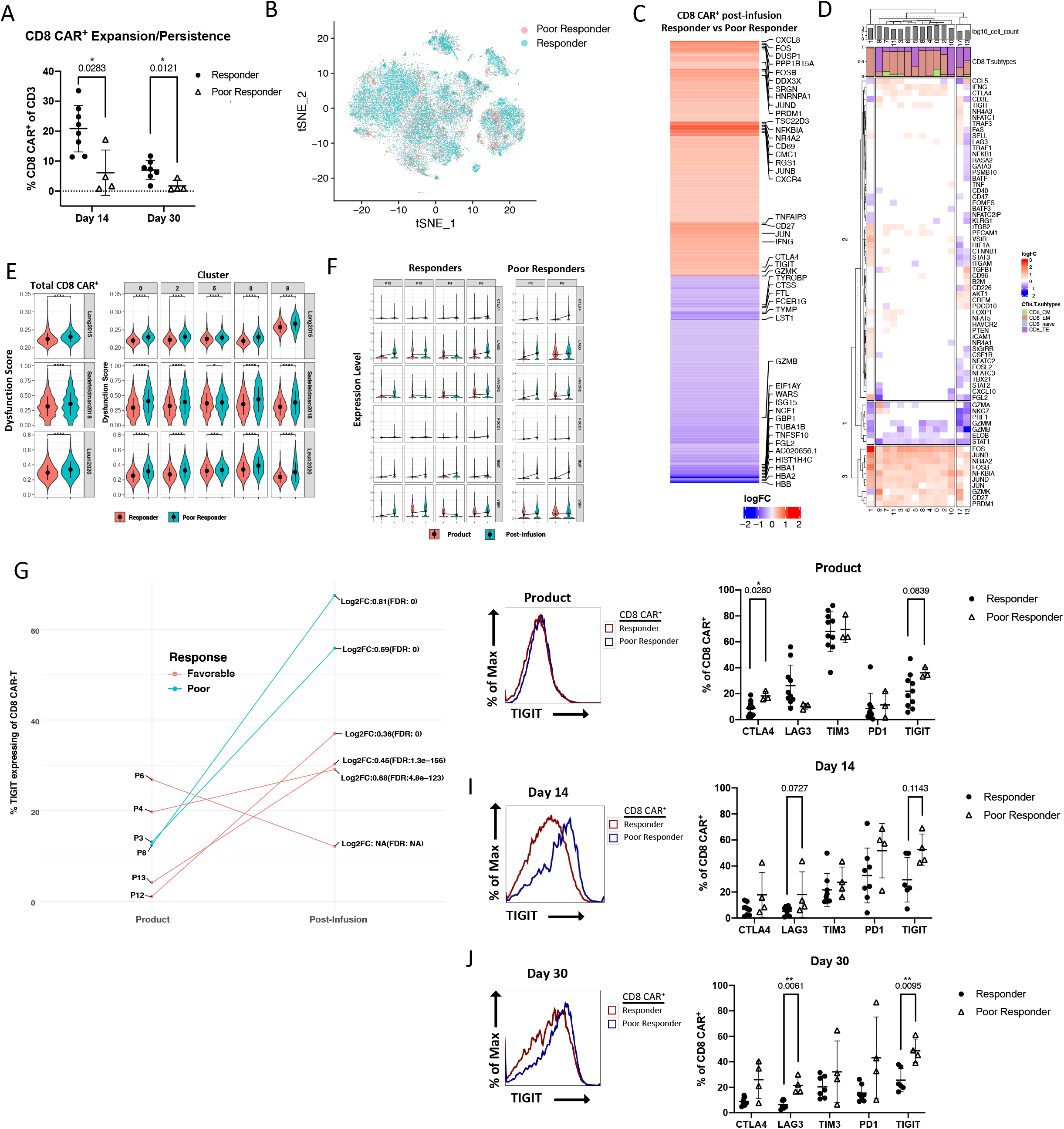
CAR-T cells of poor responders are enriched in an exhaustion-like phenotype post-infusion with high TIGIT expression. **A)** Percentage of CD8 CAR-T cells of total T cells between response groups at day 14 and day 30 as measured by flow cytometry. **B)** scRNA tSNE dimension reduction with patient response group overlay. **C)** Heatmap of differentially expressed genes (adjusted p-value < 0.05) from an immunoregulatory gene list between post-infusion CD8 CAR-T cells of response groups. Color represents log fold increase/decrease in poor responders. **D)** Heatmap of differentially expressed genes (adjusted p-value < 0.05) from an immunoregulatory gene list between post-infusion CD8 CAR-T cells of response groups within individual clusters. Color represents log fold increase/decrease in poor responders. **E)** Violin plot comparison of CAR-T cell dysfunction scores between response groups with three gene sets. Left – comparison of total CD8 CAR-T cells between response groups. Right – comparison of CD8 CAR-T cells between response groups within the most predominant clusters. **F)** Violin plots of exhaustion marker normalized RNA expression before and after infusion in individual patients. **G)** Comparison between response groups of the percentage of TIGIT expressing CD8 CAR-T cells before and after infusion. Average log fold change refers to read counts. **H-J)** Left – Histograms of fluorescence intensity of TIGIT between response groups as measured by flow cytometry. Each curve represents a concatenation of all samples with equal proportions of CD8 CAR-T cells from each sample. Right – Comparison between response groups of the percentage of CD8 CAR-T cells expressing checkpoint receptors CTLA4, LAG3, TIM3, PD1, or TIGIT.

Consistent with the transcriptional analysis of TIGIT by scRNAseq, measurement of TIGIT protein expression by flow cytometry showed a greater than 20% increase in the average percentage of TIGIT^+^ CD8 CAR-T cells in the post-infusion poor responder samples (Figure 3H-J). Interestingly, a comparison of TIGIT expression between endogenous CD8 T cells of poor and favorable response groups showed a similar trend at day 14 (p = 0.11) and day 30 (p = 0.17) (Supplemental Figure 3B). Notably, consistent differences in expression of memory markers between response groups were not apparent (Supplemental Figure 3C). Overall, our data support that poor responders in our trial to CD19 CAR-T cells have an enrichment in exhausted CD8 CAR-T cells and TIGIT is a novel prognostic marker of response.

### TIGIT expression is increased in CAR-T cells with an exhaustion phenotype

TIGIT expression has been associated with a dysfunctional T cell phenotype in chronic infection and cancer^30–32^. Nonetheless, TIGIT’s role in CAR-T cell dysfunction has not been explored. Given our previous flow cytometry analyses showed TIGIT was the checkpoint receptor most expressed after infusion and in poor responder CD8 CAR-T cells, we assessed if TIGIT can serve as a marker of CAR-T cell dysfunction. To address this question, we separated TIGIT^+^ and TIGIT^−^ CD8 CAR-T cells *in silico* and performed differential gene expression. Similar to the profile observed comparing favorable and poor responders, total TIGIT^+^ cells overexpressed genes associated with exhaustion (Figure 4A). At pre-infusion, upregulated genes included PDCD1, LAG3, EOMES, and PRDM1; downregulated genes included TCF7, SELL, and CCR7 (padj < 0.05). At post-infusion, TIGIT^+^ cells had elevated levels of exhaustion markers TOX, PD1, and GZMK. Elevated TIGIT expression was not associated with any particular cluster or cell subtype in post-infusion CD8 CAR-T cells (Figure 4C; Supplemental Figure 4A). We next applied T cell dysfunction scoring to compare TIGIT^+^ cells to TIGIT^−^ cells. TIGIT^+^ cells had higher dysfunction scores globally (p < 0.0001) and in the most predominant clusters across the three gene sets tested (Figure 4C). Furthermore, high dysfunction scores co-localized with TIGIT expression (Figure 4B;C). Accordingly, upon comparison of exhaustion marker expression in TIGIT^+^ versus TIGIT^−^ CD8 CAR-T cells across all three time points, we observed significantly increased expression of CTLA4, LAG3, and PD1 was observed with average fold changes of 1.94, 1.95, and 1.48, respectively (p < 0.05) (Figure 4D-F). Interestingly, the same trend was observed in CD8 T cells from patients that did not express the CAR, with fold increases in CTLA4 (3.04), LAG3 (2.39), PD1 (1.97), and TIM3 (1.78) (Supplemental Figure 2B).

**Figure 4.**
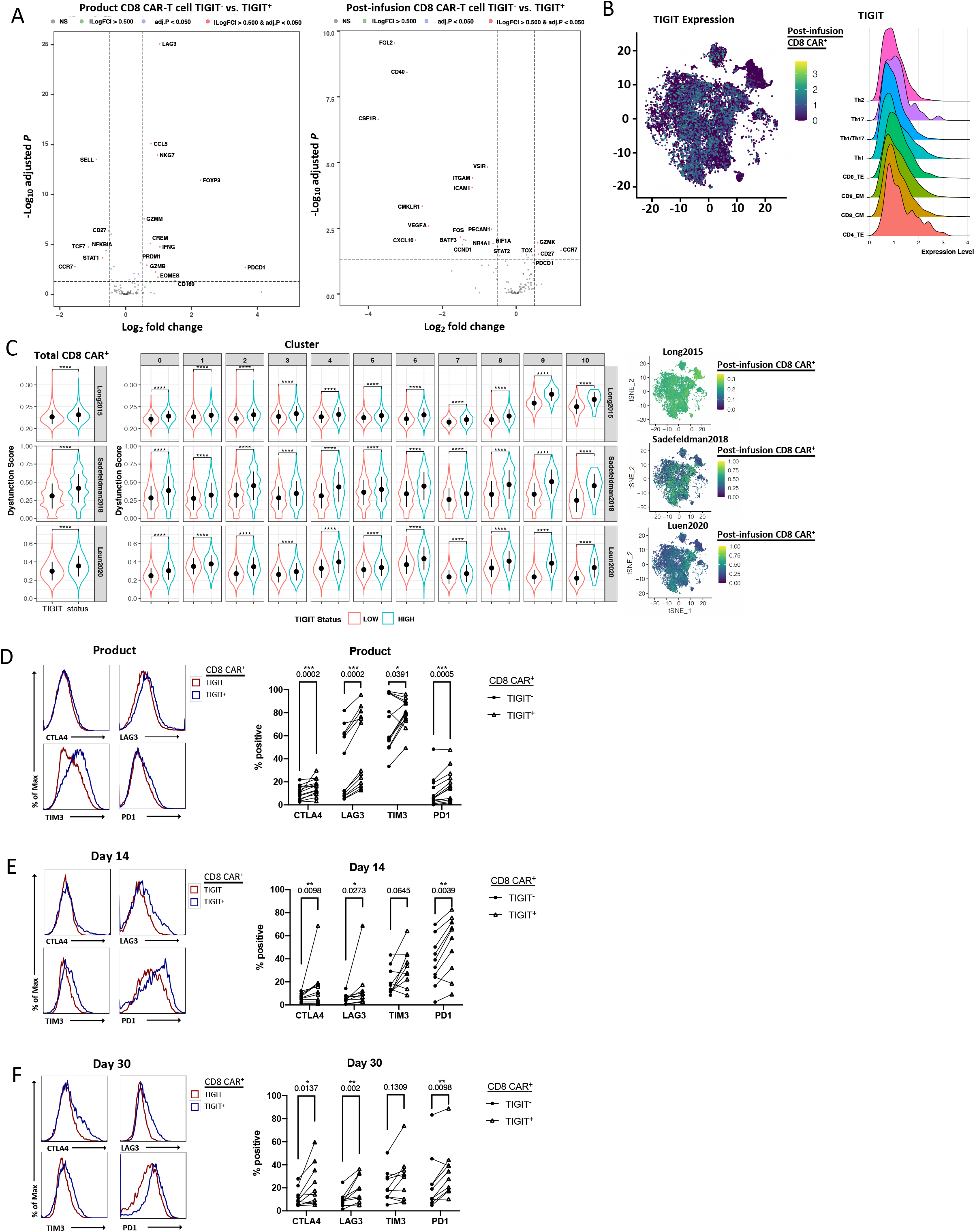
TIGIT expression is increased in CAR-T cells with an exhaustion phenotype. **A)** Volcano plot of differentially expressed genes (adjusted p-value < 0.05) between TIGIT^+^ and TIGIT^−^ CD8 CAR-T cells in pre-infusion or post-infusion samples by scRNA seq. **B)** Left - scRNA tSNE dimension reduction with TIGIT RNA expression overlay in post-infusion CD8 CAR-T cells. Right – Ridge plot of TIGIT RNA expression across cell subtype assignments. **C)** Left - Violin plot comparison of dysfunction scores between TIGIT^+^ and TIGIT^−^ cells of total CD8 CART cells or of CD8 CAR-T cells within individual clusters. Columns indicate clusters and rows indicate exhaustion gene set utilized. Right – scRNA tSNE dimension reduction plots with dysfunction scores overlaid on post-infusion CD8 CAR-T cells. Each plot corresponds to the indicated exhaustion gene set. **D-F)** Left – Histograms of fluorescence intensity of checkpoint receptor expression comparing between TIGIT^+^ and TIGIT^−^ CD8 CAR-T cells at the indicated time point as measured by flow cytometry. Each curve represents a concatenation of all samples with equal proportions of CD8 CAR-T cells from each sample. Right – Comparison of the percentage of cells expressing checkpoint receptors CTLA4, LAG3, TIM3, or PD1 between TIGIT^+^ and TIGIT^−^ CD8 CAR-T cells.

### Functional inhibition of CAR-T cells by TIGIT expression

TIGIT is thought to inhibit T cells by competing with the co-stimulatory receptor DNAM-1 in binding to their common ligands PVR and PVRL2^33, 34^. Studies also suggest that TIGIT can bind to DNAM-1 in cis, disrupting DNAM-1 homodimerization, or it may directly inhibit T cell activation by signaling through its inhibitory ITT and ITIM domains^32, 35^. In NHL, PVR expression has been reported on tumor cells and endothelial cells^30^; other contexts have reported expression on intratumoral myeloid cells^36, 37^. To determine if TIGIT competition with DNAM-1 for PVR and PVRL2 binding is a potential mechanism of CAR-T cell dysfunction, we first measured DNAM-1 expression on CAR-T cells from the clinical product and after infusion. As shown in Figure 5A, nearly all of the CD8 CAR-T cells express very high levels of DNAM-1 in the product, and upon infusion the CD8 CAR-T cells remain over 50% DNAM1^+^ in the majority of patients. We next determined if TIGIT and DNAM-1 were co-expressed, and observed the majority of TIGIT^+^ cells also express DNAM-1 at day 14 (69%) and day 30 (85%) (Figure 5B). Given the expression of PVR on target cells and neighboring cell types, TIGIT blockade in CD8 CAR^+^ cells may be a means to improve CAR T cell therapy.

**Figure 5.**
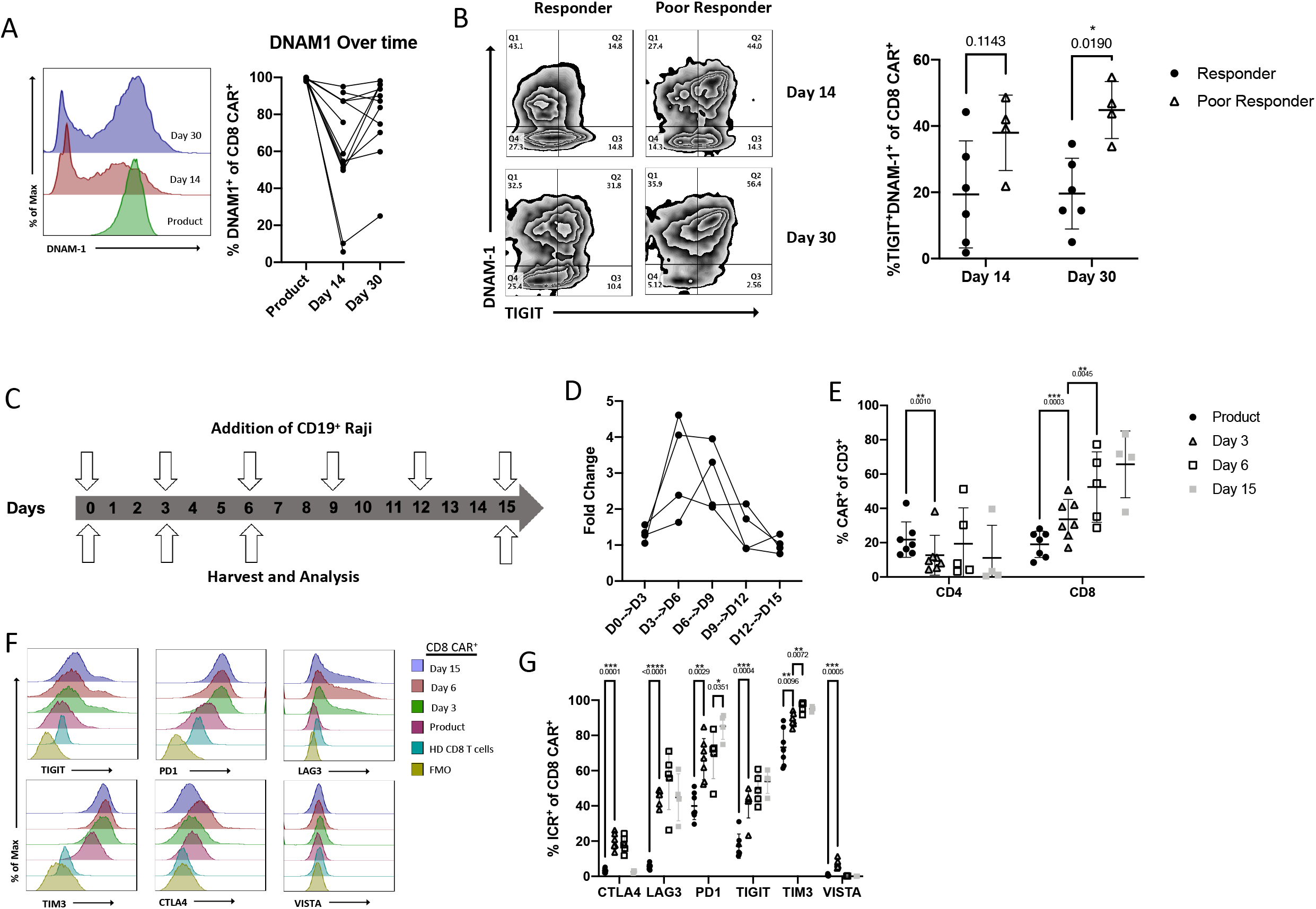
Functional inhibition of CAR-T cells by TIGIT expression. **A)** Left – Histograms of fluorescence intensity of DNAM-1 in CD8 CAR-T cells across time points as measured by flow cytometry. Each curve represents a concatenation of all samples with equal proportions of CD8 CAR-T cells from each sample. Right – Comparison between time points of the percentage of DNAM-1 expressing cells of CD8 CAR-T cells. **B)** Left - Zebra plots depicting co-expression of TIGIT and DNAM-1 in CD8 CAR-T cells as measured by flow cytometry. Indicated responder and poor responder samples are concatenations of equal proportions of CD8 CAR-T cells from each sample. Right – Percentage of TIGIT and DNAM-1 co-expression in CD8 CAR-T cells of responders and poor responders at day 14 and day 30 post-infusion. **C)** Setup of in vitro exhaustion model. CAR-T cell products were stimulated with CD19^+^ Raji every three days until day 15. At days 3, 6, and 15 aliquots were taken for flow cytometric analysis. **D)** From model shown in C). Fold change in the number of CAR-T cells over time at the indicated time points as determined by flow cytometric analysis and cell counts. **E)** Frequency of CD4 or CD8 CAR-T cells of total T cells across days 0, 3, 6, and 15 of the exhaustion model shown in C). **F)** From model shown in C), comparison of fluorescence intensity of checkpoint receptors TIGIT, PD1, LAG3, TIM3, CTLA4, and VISTA over time in CD8 CAR-T cells. Each curve represents a concatenation of all samples with equal proportions of CD8 CAR-T cells from each sample. HD represents CD8 T cells from unmanipulated healthy donor peripheral blood mononuclear cells. FMO represents fluorescence minus one controls. **G)** From model shown in C), comparison across indicated time points of the percentage of CD8 CAR-T cells expressing checkpoint receptors CTLA4, LAG3, PD1, TIGIT, TIM3, or VISTA as measured by flow cytometry.

To recapitulate the CAR-T cell exhaustion we observed *in vivo*, we designed an *in vitro* chronic stimulation model. In this model, the clinical products used to treat NHL patients were stimulated with CD19^+^ Raji lymphoma cells every three days at a 4:1 ratio of CAR-T cells to Raji cells. Pre-stimulation and at days 3, 6, and 15 an aliquot of cells was removed from culture and analyzed by flow cytometry (Figure 5C). CAR^+^ cells grew robustly until day 12 of culture whereupon they stopped proliferating and failed to clear the remaining target cells, indicative of having reached an exhausted state (Figure 5D). When comparing the proportions of CD4 and CD8 CAR-T cells, an increase in the CD8/CD4 cell ratio was observed similar to the trend found *in vivo* (Figure 5E). We next compared expression of the exhaustion markers CTLA4, LAG3, PD1, TIGIT, TIM3, and VISTA across time points. Akin to the clinical setting, we observed increased expression of TIGIT in CD8 CAR-T cells from 17% at pre-stimulation to 54% at day 15 (Figure 5F;G). In contrast to the clinical setting but consistent with CAR-signaling induced expression, we observed the elevated TIM3 levels present in the product continued to increase over time. Thus, we were able to recapitulate TIGIT induction with a chronic stimulation model of CD8 CAR-T cells.

## Discussion

Here, we endeavored to compare differences in the transcriptional and phenotypic profiles of CAR-T cells between time points and response groups. We have shown that CAR-T cells isolated from the blood of patients treated with CAR-T cell therapy are antigen experienced with the transcriptional and surface marker profiles of highly activated and differentiated T cells^18^. According to both memory marker expression and gene expression cell identification, we observed heterogeneity in the memory phenotypes of post-infusion CD8 CAR-T cells including central memory, effector memory, and terminal effector cell types. Overall, we observed a shift in the predominant profiles of CD8 CAR-T cells from CD45RA^HI^CCR7^HI^CD127^HI^CD62L^HI^CD25^HI^ to CD45RO^HI^CD28^HI^CD69^HI^CD27^HI^PD1^HI^ and CD45RA^HI^CD57^HI^CD69^HI^PD1^HI^ upon infusion. We also noted upregulation of several transcription factors thought to drive exhaustion (TOX, NR4A2) and exhaustion markers TIGIT and PD1, which were frequently co-expressed. These results are in agreement with a prior scRNA sequencing study of 4 patients with complete response to CAR T cell therapy that showed greater than 60% of cells co-expressed 4 or more checkpoint receptors at 3 time points after days 7-14^17, 18^. Overall, our data supports the hypothesis that chronic stimulation drives exhaustion in post-infusion CD8 CAR-T cells. In comparison of CAR-T cells and endogenous T cells, we noted by flow cytometry that TIM3, TIGIT, and CD27 expression differentiated the groups best, which may be beneficial in future applications.

We further demonstrate that CD8 CAR-T cells from patients with progressive or stable disease are enriched with an exhaustion profile. We noted that among the tested exhaustion markers, TIGIT was the most differentially expressed among response groups. The general lack of differences in the relative frequencies in clusters between response groups and the consistent enrichment of the exhaustion profiles in poor responders across clusters and cell types indicates differences in gene expression are subtle compared to defining characteristics such as proliferation status or cell subtype. Differential gene expression revealed overexpression of transcription factors including PRDM1, NR4A2, NFKBIA, and AP-1 family members, most of which have been implicated in driving T cell exhaustion^38, 39^. While historically AP-1 family member expression has been associated with greater anti-tumor function, a recent paper highlighted a similar profile in exhausted CAR-T cells which was attributed to decreased c-FOS and c-Jun complex formation that could be remediated with c-Jun overexpression^38^. Of note, although we did not observe significant differential expression of TOX/TOX2, two known drivers of T cell exhaustion, between response groups, differential expression of TOX between TIGIT^+^ and TIGIT^−^ cells implicated TOX is associated with increased TIGIT expression. Further work will need to be done to determine the relevance of TOX and the other noted transcription factors associated with exhaustion in 4-1BB.CAR-T cells.

As TIGIT expression was dysregulated in CAR-T cells, we surveyed the phenotype of TIGIT^+^ cells to determine its relevance as a biomarker or driver of response. CD8 TIGIT^+^ CAR-T cells had greater dysfunctional scores compared to TIGIT^−^ cells, upregulated many of the same genes differentially expressed between response groups, and had higher surface expression of exhaustion markers, particularly PD1. Endogenous TIGIT^+^ CD8 T cells also had increased exhaustion marker expression, suggesting negative selection of this population may improve CAR T cell therapy. Mechanistically, TIGIT is thought to drive dysfunction in CAR-T cells by competing with DNAM-1 for binding the ligands PVR and PVRL2 expressed on the endothelium, surrounding immune cells, and tumor tissue^30^. Importantly, TIGIT and PD1 are highly expressed on endogenous intratumoral T cells of NHL patients, and in CD4 and CD8 T cells TIGIT^+^PD1^+^ cells produced the least amount of IL-2, IFNγ, and TNF-α of the four possible combinations^30^. We likewise observed in an *in vitro* exhaustion model we developed that TIGIT^+^ CAR^+^ cells killed tumor cells less efficiently and produced less effector molecules. Furthermore, TIGIT antibody blockade could prevent inhibition by TIGIT. These findings warrant further investigation of the potential of TIGIT blockade or TIGIT and PD1 co-blockade to be used in combination with CAR T cell therapy.

## Materials and methods

### Cohort description

All patients failed at least 2 previous lines of therapy and were recruited to the study in accordance with eligibility criteria described in clinicaltrials.gov (Identifier: NCT03434769; IND 17932). Leukapheresis products were obtained from NHL patients receiving CAR-T cell therapy at University Hospitals Seidman Cancer Center under a phase I/II study and utilized within 24 hours of draw. The study was approved by the institutional review board and all patients gave written informed consent.

### CAR-T cell manufacture

CAR T cell manufacture was automated with the use of the CliniMACS Prodigy® device using the TCT software program and TS520 tubing set (Miltenyi Biotec, Bergisch Gladbach, Germany). The instrument setup and technical protocol were described by Zhu *et al.^40^.* The clinical-grade reagents applied in this process were CliniMACS Buffer, TexMACS Media, CliniMACS CD4 reagent, CliniMACS CD8 reagent, TransAct, and the cytokines IL-7 and IL-15 (Miltenyi Biotec, Bergisch Gladbach, Germany). Peripheral blood apheresis products were loaded into the machine and CD4 and CD8 T cells were isolated using CliniMACS CD4 reagent and CliniMACS CD8 reagent according to the manufacturer’s instructions. The isolated T cells were then stimulated with IL-7 and IL-15 (Miltenyi Biotec, Bergisch Gladbach, Germany) at a concentration of 25μg/2L bag of TexMACS media with 3% human AB serum (Innovative Research, Novi, MI, USA). Human AB serum was removed after the 6^th^ day of culture. The viability, purity, and potency of the products were confirmed as previously described^24^. This process was performed at the Cellular Therapy Lab of University Hospitals Cleveland Medical Center Seidman Cancer Center/Case Western Reserve University Center for Regenerative Medicine.

### Lentiviral vector

The 4-1BB.CAR construct applied in the clinical trial was developed by Lentigen, a Miltenyi Biotec company (Gaithersburg, MD, USA). The vector is composed of the FMC63 scFv, a CD8-derived hinge region, TNFRSF19-derived transmembrane domain, CD3ζ intracellular domain, and 4-1BB co-stimulatory domain.

### Single cell RNA sequencing and feature barcoding library preparation

Cryopreserved apheresis products were thawed and pre-labelled with antibodies for flow cytometry (described below) and a panel of TotalSeq™-B antibodies (described below) from Biolegend (San Diego, CA, USA). The panel of TotalSeq™-B antibodies included *CD127, CD197, CD25, CD279, CD28, CD4, CD45RA, CD45RO, CD57, CD62L, CD69,* and *CD8*. Live CD3^+^CAR^+^ T cells were sorted by FACS and preparation of single cell and TotalSeq™-B libraries was performed utilizing the 10x Genomics Chromium Single Cell 3’ Reagent kits with feature barcoding technology for cell surface protein (v3) (Pleasanton, CA, USA) according to the manufacturer’s instructions. Libraries were sequenced by Psomagen, Inc (Rockville, MD, USA).

### Quality control of raw 10x scRNA sequencing data

A total of 27 CAR-T samples were sequenced. For each sequenced scRNA-Seq pool, Cell Ranger (v3.1.0) from 10x Genomics (Pleasanton, CA, USA) was used to process, align, and summarize unique molecular identifier (UMI) counts against hg38 human reference genome. For the 16 CAR-T samples, 12 T-cell surface proteins (*CD69*, *CD62L*, *CD197*, *CD25*, *CD279*, *CD45RA*, *CD127*, *CD4*, *CD45RO*, and *CD8A*) are barcoded to each cell by TotalSeq™-B antibody library (Biolegend, San Diego, CA, USA). Based on each sample mRNA assay UMI metrics, cells with too low or high UMI counts or the number of genes (i.e., 2.5 x standard deviation) were filtered out. Cells with a mitochondrial UMI count proportion higher than 15% were removed. Cells with a ratio of number of genes covered to UMI counts less than 0.1 were also filtered. Doublets as annotated by *Scrublet* v.0.2.1^41^, were also removed. After the comprehensive quality control procedure, we only retained 24 CAR-T samples since for 3 samples (patient 5 day 14, patient 5 day 30, patient 12 day 30) either the number of read counts or the number of genes was too small compared to the other samples.

### *In silico* cell type prediction

CAR-T cell subtypes were assigned by S*ingleR*^21^ (v1.0.6). Gene expression profile was compared with the RNA-seq transcriptome profile of 29 immune cell types^22^. The best-predicted cell type was considered. CAR-T cells were grouped into CD4 and CD8 T cells. CD4^+^ T cells were annotated as T follicular helper cells (Tfh), regulatory T cells (Tregs), Th1, Th1/Th17, Th17, Th2, naive, or terminal effector. CD8^+^ T cells were annotated as naive, central memory, effector memory, or terminal effector.

### 10X Genomics scRNA sequencing data analysis and adjusting batch effects

The 24 CAR-T samples from 13 patients includes a total of 94K high-quality cells with an average of 3,917 cells per sample. The samples were merged and the *Seurat* R package^42^ (v3.2.3) was used to normalize expression values for total UMI counts per cell. Two thousand highly variable genes were identified by fitting the mean-variance relationship. Cell cycle was inferred by a function, *CellCycleScoring*, from the *Seurat* package using 43 genes for the S state and 54 genes for the G2M state, respectively. The merged samples batch effect signal was corrected using the *Harmony* (v1.0) algorithm^43^, and both mitochondrial gene expression and cell cycle annotation were regressed out. The aligned cells were then clustered using the *Louvain* algorithm for modularity optimization using the *kNN* (k nearest neighbors) graph as input. Cell clusters were visualized using the *tSNE* algorithm^44^ with a dimension reduction input from Harmony.

### Marker gene detection and differential expression analysis

For each identified cluster, we compared the cells within the clusters versus all other cells using R packages *Seurat* and *MAST*^45^ (v1.16.0) for statistical testing to identify all marker genes expressed distinctly compared to the other clusters. Only differentially-expressed genes of significance less than 5% FDR were retained. Without loss of the generality, the same DEG testing is applied between the CAR-T product and post-infusion CAR-T cells, or between the cells belonging to the patients with a favorable or poor outcome within either the same cluster or T cell subtype.

### Immune regulated genes

In this study, we focused on a customized list of 106 genes for discussion and visualization. The genes are either directly or indirectly known to be associated with human immunology^46^.

### CD8^+^ T cell dysfunctional score

CD8^+^ T cell dysfunctional scores were calculated at each cell. We computed the AUC score of a gene set showing a CD8^+^ T cell dysfunction phenotype using *AUCell_calcAUC* from R package *AUCell^47^* (v1.8.0). Three signature gene sets are used for the prediction. The first gene set includes *LAG3, PDCD1, HAVCR2, TIGIT, CD38,* and *ENTPD1^28^*. The second gene set includes a total of 22 genes: *LAYN, ITGAE, PDCD1, CTLA4, HAVCR2, LAG3, TIGIT, CXCL13, CD38, ENTPD1, CDK1, HSPH1, CCCNB1, HSPB1, MKI67, DK4, GZMB, TOX, IFNG, MIR155HG, TNFRSF9*, and *RB1^29^*. The third signature gene set was sorted by the log fold change between the controls and experiment sample group and the top 2,000 positive log fold change genes were selected for AUC calculation^1^.

### PBMC scRNA-Seq analysis

A total of 15 PBMC scRNA-Seq samples were sequenced. These samples were all obtained after CAR-T cell infusion from the same patients that participated in the CAR T cell study. After the same quality control that was done in the CAR T samples was applied, we retained a total of 75,324 PBMC cells. Similarly, each cell type was annotated by R package *SingleR* with the PBMC cell type mRNA reference profile^22^. As a result, each cell was annotated with either γδ T cells, monocytes, natural killer (NK) cells, CD8^+^ T, CD4^+^ T, dendritic, progenitor, B cells, neutrophils, or basophils.

### Flow cytometry

Extracellular staining for flow cytometry was performed by incubating titrated amounts of fluorescent-labelled antibodies or viability dye for 15 minutes at room temperature. Secondary staining for biotin-streptavidin conjugates was performed with a 30-minute incubation at room temperature. Acquisition was performed with a BD ARIA flow cytometer (BD Biosciences, Franklin Lakes, NJ, USA). See table for complete list of flow cytometric reagents (Supplemental Table 4).

### Flow cytometric analyses and statistics

Flow cytometric gating was based on fluorescence minus one controls in cases where bimodal distribution was not apparent. To generate histogram comparisons of fluorescence intensity across samples run with the same cytometer settings on different days, we first performed batch correction with the function *SwiftReg^48^* on *MATLAB_R2020b* (The MathWorks, Inc., Natick, MA, USA). Comparisons were then performed with concatenations of all samples containing equal proportions of the cell type of interest. For comparison of two groups without matching, a Mann-Whitney test was performed. Comparisons of two groups with matching samples were performed with Wilcoxon matched-pairs signed rank tests. For comparisons of three or more groups, a Friedman test was performed. For paired comparisons across three or more groups, a mixed-effects analysis was performed. For all plots derived from flow cytometric data, no correction was made for multiple comparisons and all comparisons made in the statistical analysis are displayed. All statistical analyses were done using *Prism 9.0.2* (GraphPad Software, San Diego, CA, USA). Statistical significance was provided by asterisk, and the number of asterisks shown correspond to p-values less than 0.05, 0.01, 0.001, or 0.0001, respectively.

### Exhaustion assay

CAR-T cells were thawed and rested overnight in 5ng/mL IL-15 and 10ng/mL IL-17 in cRPMI. The next day, CAR products were either FACS sorted with anti-FMC63-FITC to obtain a pure CAR^+^ population or immediately added to culture with Raji cell line at a 4:1 CAR-T cell to Raji cell ratio in 30U/mL IL-2. Every three days, CAR-T cells were counted and the effector to target ratio restored with fresh cRPMI and 30U/mL IL-2. Aliquots were taken at the indicated days for flow cytometric analysis.

## Data Sharing Statement

Raw sequencing data will be uploaded to The European Genome-phenome Archive (EGA) and accessible to the public when the manuscript is published.

## Acknowledgements

This research was supported by the Case Comprehensive Cancer Center (P30CA043703) including the Hematopoietic Biorepository and Cellular Therapy Shared Resource and the Cytometry and Microscopy Shared Resource as well as the NIH grant T32AI089474.

## Authorship

D.N.W conceived of the project. D.N.W and T.H. supervised the study, assisted in data analysis, and funded the study. Z.J., K.G. and R.S. performed experimental work. C.H., R.S. and Z.J. performed bioinformatics data analysis. P.F.C. and M.L. led the clinical trial and performed sample banking. J.R. helped oversee sample banking. Z.J. wrote the manuscript. All authors contributed to editing the manuscript.

## References

1. Long, AH, WM Haso, JF Shern, KM Wanhainen, M Murgai, M Ingaramo, JP Smith, AJ Walker, ME Kohler, VR Venkateshwara, RN Kaplan, GH Patterson, TJ Fry, RJ Orentas, and CL Mackall, 4-1BB costimulation ameliorates T cell exhaustion induced by tonic signaling of chimeric antigen receptors. Nat Med. 2015;21(6):581–90.

2. Ying, Z, T He, X Wang, W Zheng, N Lin, M Tu, Y Xie, L Ping, C Zhang, W Liu, L Deng, F Qi, Y Ding, XA Lu, Y Song, and J Zhu, Parallel Comparison of 4-1BB or CD28 Co-stimulated CD19-Targeted CAR-T Cells for B Cell Non-Hodgkin’s Lymphoma. Mol Ther Oncolytics. 2019;15:60–68.

3. Weinkove, R, P George, N Dasyam, and AD McLellan, Selecting costimulatory domains for chimeric antigen receptors: functional and clinical considerations. Clin Transl Immunology. 2019;8(5):e1049.

4. Ansell, SM, Non-Hodgkin Lymphoma: Diagnosis and Treatment. Mayo Clin Proc. 2015;90(8):1152–63.

5. Gribben, JG and S O’Brien, Update on therapy of chronic lymphocytic leukemia. J Clin Oncol. 2011;29(5):544–50.

6. Cortelazzo, S, M Ponzoni, AJ Ferreri, and M Dreyling, Mantle cell lymphoma. Crit Rev Oncol Hematol. 2012;82(1):78–101.

7. Crump, M, SS Neelapu, U Farooq, E Van Den Neste, J Kuruvilla, J Westin, BK Link, A Hay, JR Cerhan, L Zhu, S Boussetta, L Feng, MJ Maurer, L Navale, J Wiezorek, WY Go, and C Gisselbrecht, Outcomes in refractory diffuse large B-cell lymphoma: results from the international SCHOLAR-1 study. Blood. 2017;130(16):1800–1808.

8. Neelapu, SS, FL Locke, NL Bartlett, LJ Lekakis, DB Miklos, CA Jacobson, I Braunschweig, OO Oluwole, T Siddiqi, Y Lin, JM Timmerman, PJ Stiff, JW Friedberg, IW Flinn, A Goy, BT Hill, MR Smith, A Deol, U Farooq, P McSweeney, J Munoz, I Avivi, JE Castro, JR Westin, JC Chavez, A Ghobadi, KV Komanduri, R Levy, ED Jacobsen, TE Witzig, P Reagan, A Bot, J Rossi, L Navale, Y Jiang, J Aycock, M Elias, D Chang, J Wiezorek, and WY Go, Axicabtagene Ciloleucel CAR T-Cell Therapy in Refractory Large B-Cell Lymphoma. N Engl J Med. 2017;377(26):2531–2544.

9. Kersten, MJ, AM Spanjaart, and C Thieblemont, CD19-directed CAR T-cell therapy in B-cell NHL. Curr Opin Oncol. 2020;32(5):408–417.

10. Abramson, JS, M Lunning, and ML Palomba, Chimeric Antigen Receptor T-Cell Therapies for Aggressive B-Cell Lymphomas: Current and Future State of the Art. Am Soc Clin Oncol Educ Book. 2019;39:446–453.

11. Hunter, BD, M Rogalski, and CA Jacobson, Chimeric antigen receptor T-cell therapy for the treatment of aggressive B-cell non-Hodgkin lymphomas: efficacy, toxicity, and comparative chimeric antigen receptor products. Expert Opin Biol Ther. 2019;19(11):1157–1164.

12. Jacoby, E, SA Shahani, and NN Shah, Updates on CAR T-cell therapy in B-cell malignancies. Immunol Rev. 2019;290(1):39–59.

13. Chavez, JC, C Bachmeier, and MA Kharfan-Dabaja, CAR T-cell therapy for B-cell lymphomas: clinical trial results of available products. Ther Adv Hematol. 2019;10:2040620719841581.

14. Chong, EA, M Ruella, SJ Schuster, and P Lymphoma Program Investigators at the University of, Five-Year Outcomes for Refractory B-Cell Lymphomas with CAR T-Cell Therapy. N Engl J Med. 2021;384(7):673–674.

15. Shah, NN and TJ Fry, Mechanisms of resistance to CAR T cell therapy. Nat Rev Clin Oncol. 2019;16(6):372–385.

16. Deng, Q, G Han, N Puebla-Osorio, MCJ Ma, P Strati, B Chasen, E Dai, M Dang, N Jain, H Yang, Y Wang, S Zhang, R Wang, R Chen, J Showell, S Ghosh, S Patchva, Q Zhang, R Sun, F Hagemeister, L Fayad, F Samaniego, HC Lee, LJ Nastoupil, N Fowler, R Eric Davis, J Westin, SS Neelapu, L Wang, and MR Green, Characteristics of anti-CD19 CAR T cell infusion products associated with efficacy and toxicity in patients with large B cell lymphomas. Nat Med. 2020;26(12):1878–1887.

17. Fraietta, JA, SF Lacey, EJ Orlando, I Pruteanu-Malinici, M Gohil, S Lundh, AC Boesteanu, Y Wang, RS O’Connor, WT Hwang, E Pequignot, DE Ambrose, C Zhang, N Wilcox, F Bedoya, C Dorfmeier, F Chen, L Tian, H Parakandi, M Gupta, RM Young, FB Johnson, I Kulikovskaya, L Liu, J Xu, SH Kassim, MM Davis, BL Levine, NV Frey, DL Siegel, AC Huang, EJ Wherry, H Bitter, JL Brogdon, DL Porter, CH June, and JJ Melenhorst, Determinants of response and resistance to CD19 chimeric antigen receptor (CAR) T cell therapy of chronic lymphocytic leukemia. Nat Med. 2018;24(5):563–571.

18. Sheih, A, V Voillet, LA Hanafi, HA DeBerg, M Yajima, R Hawkins, V Gersuk, SR Riddell, DG Maloney, ME Wohlfahrt, D Pande, MR Enstrom, HP Kiem, JE Adair, R Gottardo, PS Linsley, and CJ Turtle, Clonal kinetics and single-cell transcriptional profiling of CAR-T cells in patients undergoing CD19 CAR-T immunotherapy. Nat Commun. 2020;11(1):219.

19. Stoeckius, M, C Hafemeister, W Stephenson, B Houck-Loomis, PK Chattopadhyay, H Swerdlow, R Satija, and P Smibert, Simultaneous epitope and transcriptome measurement in single cells. Nat Methods. 2017;14(9):865–868.

20. Lee, MS, K Hanspers, CS Barker, AP Korn, and JM McCune, Gene expression profiles during human CD4+ T cell differentiation. Int Immunol. 2004;16(8):1109–24.

21. Aran, D, AP Looney, L Liu, E Wu, V Fong, A Hsu, S Chak, RP Naikawadi, PJ Wolters, AR Abate, AJ Butte, and M Bhattacharya, Reference-based analysis of lung single-cell sequencing reveals a transitional profibrotic macrophage. Nat Immunol. 2019;20(2):163–172.

22. Monaco, G, B Lee, W Xu, S Mustafah, YY Hwang, C Carre, N Burdin, L Visan, M Ceccarelli, M Poidinger, A Zippelius, J Pedro de Magalhaes, and A Larbi, RNA-Seq Signatures Normalized by mRNA Abundance Allow Absolute Deconvolution of Human Immune Cell Types. Cell Rep. 2019;26(6):1627–1640 e7.

23. Turtle, CJ, LA Hanafi, C Berger, TA Gooley, S Cherian, M Hudecek, D Sommermeyer, K Melville, B Pender, TM Budiarto, E Robinson, NN Steevens, C Chaney, L Soma, X Chen, C Yeung, B Wood, D Li, J Cao, S Heimfeld, MC Jensen, SR Riddell, and DG Maloney, CD19 CAR-T cells of defined CD4+:CD8+ composition in adult B cell ALL patients. J Clin Invest. 2016;126(6):2123–38.

24. Jackson, Z, A Roe, AA Sharma, F Lopes, A Talla, S Kleinsorge-Block, K Zamborsky, J Schiavone, S Manjappa, R Schauner, G Lee, R Liu, PF Caimi, Y Xiong, W Krueger, A Worden, M Kadan, D Schneider, R Orentas, B Dropulic, RP Sekaly, M de Lima, DN Wald, and JS Reese, Automated Manufacture of Autologous CD19 CAR-T Cells for Treatment of Non-hodgkin Lymphoma. Front Immunol. 2020;11:1941.

25. Martin, MD and VP Badovinac, Defining Memory CD8 T Cell. Front Immunol. 2018;9:2692.

26. Strioga, M, V Pasukoniene, and D Characiejus, CD8+ CD28- and CD8+ CD57+ T cells and their role in health and disease. Immunology. 2011;134(1):17–32.

27. Hendriks, J, LA Gravestein, K Tesselaar, RA van Lier, TN Schumacher, and J Borst, CD27 is required for generation and long-term maintenance of T cell immunity. Nat Immunol. 2000;1(5):433–40.

28. Sade-Feldman, M, K Yizhak, SL Bjorgaard, JP Ray, CG de Boer, RW Jenkins, DJ Lieb, JH Chen, DT Frederick, M Barzily-Rokni, SS Freeman, A Reuben, PJ Hoover, AC Villani, E Ivanova, A Portell, PH Lizotte, AR Aref, JP Eliane, MR Hammond, H Vitzthum, SM Blackmon, B Li, V Gopalakrishnan, SM Reddy, ZA Cooper, CP Paweletz, DA Barbie, A Stemmer-Rachamimov, KT Flaherty, JA Wargo, GM Boland, RJ Sullivan, G Getz, and N Hacohen, Defining T Cell States Associated with Response to Checkpoint Immunotherapy in Melanoma. Cell. 2019;176(1–2):404.

29. van der Leun, AM, DS Thommen, and TN Schumacher, CD8(+) T cell states in human cancer: insights from single-cell analysis. Nat Rev Cancer. 2020;20(4):218–232.

30. Josefsson, SE, K Beiske, YN Blaker, MS Forsund, H Holte, B Ostenstad, E Kimby, H Koksal, S Walchli, B Bai, EB Smeland, R Levy, A Kolstad, K Huse, and JH Myklebust, TIGIT and PD-1 Mark Intratumoral T Cells with Reduced Effector Function in B-cell Non-Hodgkin Lymphoma. Cancer Immunol Res. 2019;7(3):355–362.

31. Chew, GM, T Fujita, GM Webb, BJ Burwitz, HL Wu, JS Reed, KB Hammond, KL Clayton, N Ishii, M Abdel-Mohsen, T Liegler, BI Mitchell, FM Hecht, M Ostrowski, CM Shikuma, SG Hansen, M Maurer, AJ Korman, SG Deeks, JB Sacha, and LC Ndhlovu, TIGIT Marks Exhausted T Cells, Correlates with Disease Progression, and Serves as a Target for Immune Restoration in HIV and SIV Infection. PLoS Pathog. 2016;12(1):e1005349.

32. Johnston, RJ, L Comps-Agrar, J Hackney, X Yu, M Huseni, Y Yang, S Park, V Javinal, H Chiu, B Irving, DL Eaton, and JL Grogan, The immunoreceptor TIGIT regulates antitumor and antiviral CD8(+) T cell effector function. Cancer Cell. 2014;26(6):923–937.

33. Levin, SD, DW Taft, CS Brandt, C Bucher, ED Howard, EM Chadwick, J Johnston, A Hammond, K Bontadelli, D Ardourel, L Hebb, A Wolf, TR Bukowski, MW Rixon, JL Kuijper, CD Ostrander, JW West, J Bilsborough, B Fox, Z Gao, W Xu, F Ramsdell, BR Blazar, and KE Lewis, Vstm3 is a member of the CD28 family and an important modulator of T-cell function. Eur J Immunol. 2011;41(4):902–15.

34. Lozano, E, M Dominguez-Villar, V Kuchroo, and DA Hafler, The TIGIT/CD226 axis regulates human T cell function. J Immunol. 2012;188(8):3869–75.

35. Joller, N, JP Hafler, B Brynedal, N Kassam, S Spoerl, SD Levin, AH Sharpe, and VK Kuchroo, Cutting edge: TIGIT has T cell-intrinsic inhibitory functions. J Immunol. 2011;186(3):1338–42.

36. Gao, J, Q Zheng, N Xin, W Wang, and C Zhao, CD155, an onco-immunologic molecule in human tumors. Cancer Sci. 2017;108(10):1934–1938.

37. Zhu, Y, A Paniccia, AC Schulick, W Chen, MR Koenig, JT Byers, S Yao, S Bevers, and BH Edil, Identification of CD112R as a novel checkpoint for human T cells. J Exp Med. 2016;213(2):167–76.

38. Lynn, RC, EW Weber, E Sotillo, D Gennert, P Xu, Z Good, H Anbunathan, J Lattin, R Jones, V Tieu, S Nagaraja, J Granja, CFA de Bourcy, R Majzner, AT Satpathy, SR Quake, M Monje, HY Chang, and CL Mackall, c-Jun overexpression in CAR T cells induces exhaustion resistance. Nature. 2019;576(7786):293–300.

39. Seo, H, J Chen, E Gonzalez-Avalos, D Samaniego-Castruita, A Das, YH Wang, IF Lopez-Moyado, RO Georges, W Zhang, A Onodera, CJ Wu, LF Lu, PG Hogan, A Bhandoola, and A Rao, TOX and TOX2 transcription factors cooperate with NR4A transcription factors to impose CD8(+) T cell exhaustion. Proc Natl Acad Sci U S A. 2019;116(25):12410–12415.

40. Zhu, F, N Shah, H Xu, D Schneider, R Orentas, B Dropulic, P Hari, and CA Keever-Taylor, Closed-system manufacturing of CD19 and dual-targeted CD20/19 chimeric antigen receptor T cells using the CliniMACS Prodigy device at an academic medical center. Cytotherapy. 2018;20(3):394–406.

41. Wolock, SL, R Lopez, and AM Klein, Scrublet: Computational Identification of Cell Doublets in Single-Cell Transcriptomic Data. Cell Syst. 2019;8(4):281–291 e9.

42. Butler, A, P Hoffman, P Smibert, E Papalexi, and R Satija, Integrating single-cell transcriptomic data across different conditions, technologies, and species. Nat Biotechnol. 2018;36(5):411–420.

43. Korsunsky, I, N Millard, J Fan, K Slowikowski, F Zhang, K Wei, Y Baglaenko, M Brenner, PR Loh, and S Raychaudhuri, Fast, sensitive and accurate integration of single-cell data with Harmony. Nat Methods. 2019;16(12):1289–1296.

44. van der Maaten, LJPGEH, Visualizing High-Dimensional Data Using t-SNE. Journal of Machine Learning Research. 2008;9:2579––2605.

45. Finak, G, A McDavid, M Yajima, J Deng, V Gersuk, AK Shalek, CK Slichter, HW Miller, MJ McElrath, M Prlic, PS Linsley, and R Gottardo, MAST: a flexible statistical framework for assessing transcriptional changes and characterizing heterogeneity in single-cell RNA sequencing data. Genome Biol. 2015;16:278.

46. Bhattacharya, S, P Dunn, CG Thomas, B Smith, H Schaefer, J Chen, Z Hu, KA Zalocusky, RD Shankar, SS Shen-Orr, E Thomson, J Wiser, and AJ Butte, ImmPort, toward repurposing of open access immunological assay data for translational and clinical research. Sci Data. 2018;5:180015.

47. Aibar, S, CB Gonzalez-Blas, T Moerman, VA Huynh-Thu, H Imrichova, G Hulselmans, F Rambow, JC Marine, P Geurts, J Aerts, J van den Oord, ZK Atak, J Wouters, and S Aerts, SCENIC: single-cell regulatory network inference and clustering. Nat Methods. 2017;14(11):1083–1086.

48. Rebhahn, JA, SA Quataert, G Sharma, and TR Mosmann, SwiftReg cluster registration automatically reduces flow cytometry data variability including batch effects. Commun Biol. 2020;3(1):218.

