## Supplemental Material for "Sequential single cell transcriptional and protein marker profiling reveals TIGIT as a marker of CD19 CAR-T cell dysfunction in patients with non-Hodgkin’s lymphoma"

**Supplemental Tables**

**Supplemental Table 1 – Sample Info**

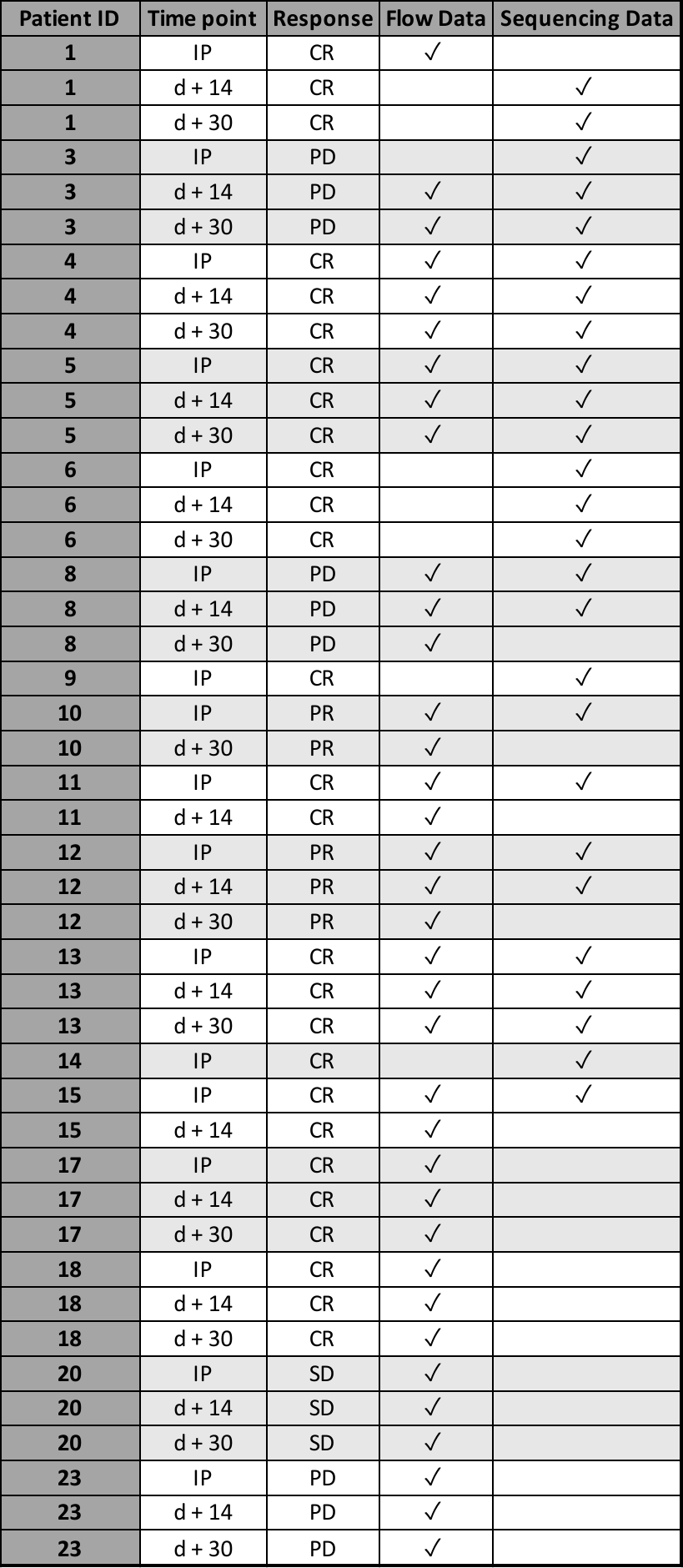

**Supplemental Table 2 – Clinical Info**

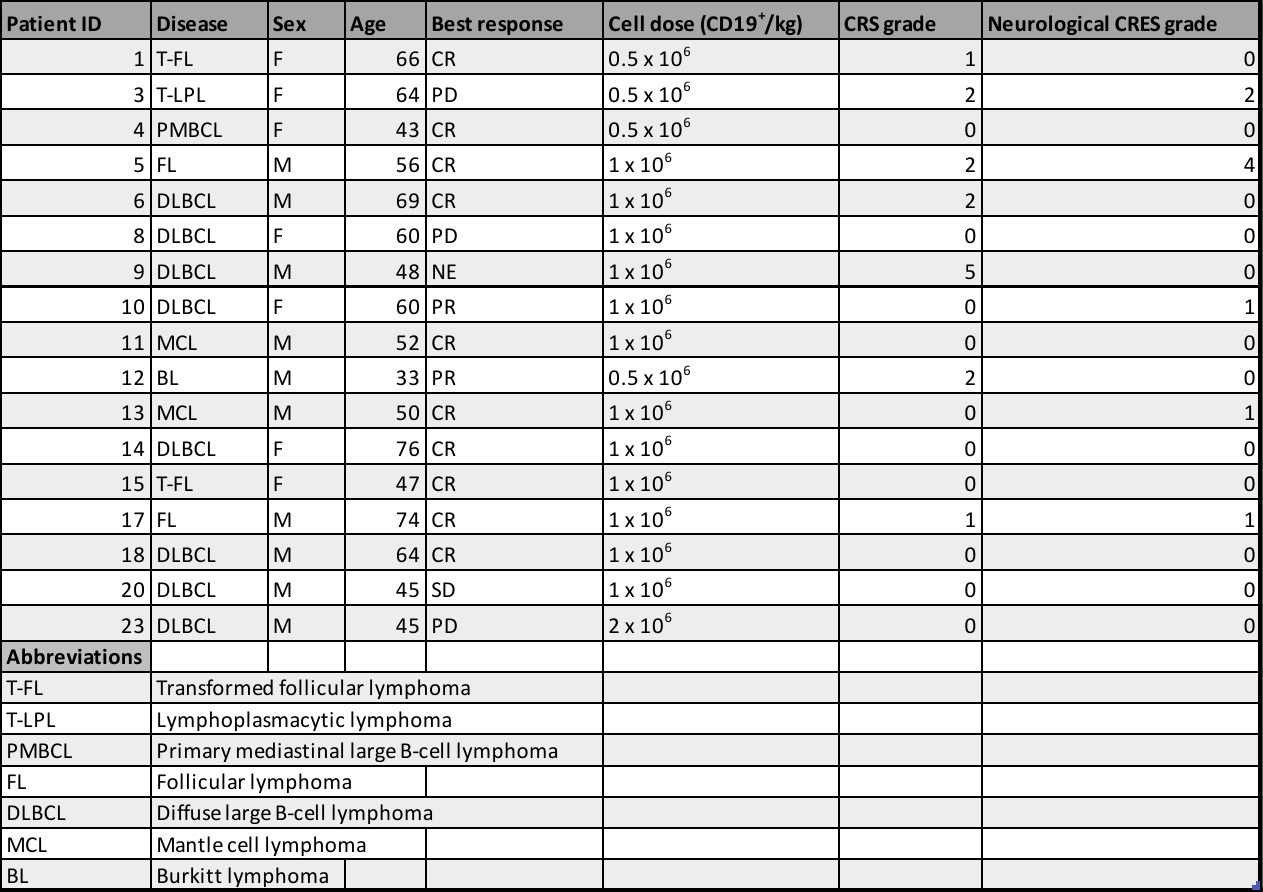

**Supplemental Table 3 – Quality Control Metrics**

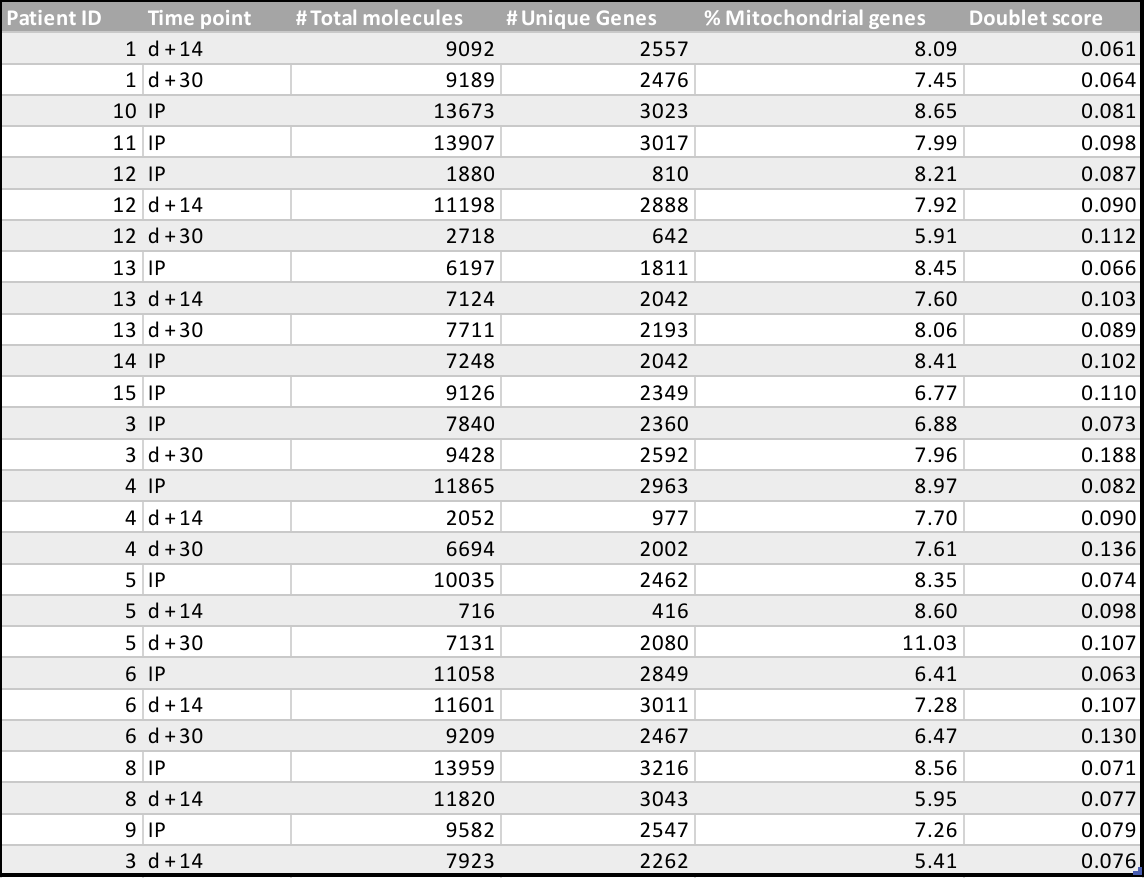

**Supplemental Table 4 – Reagents List**

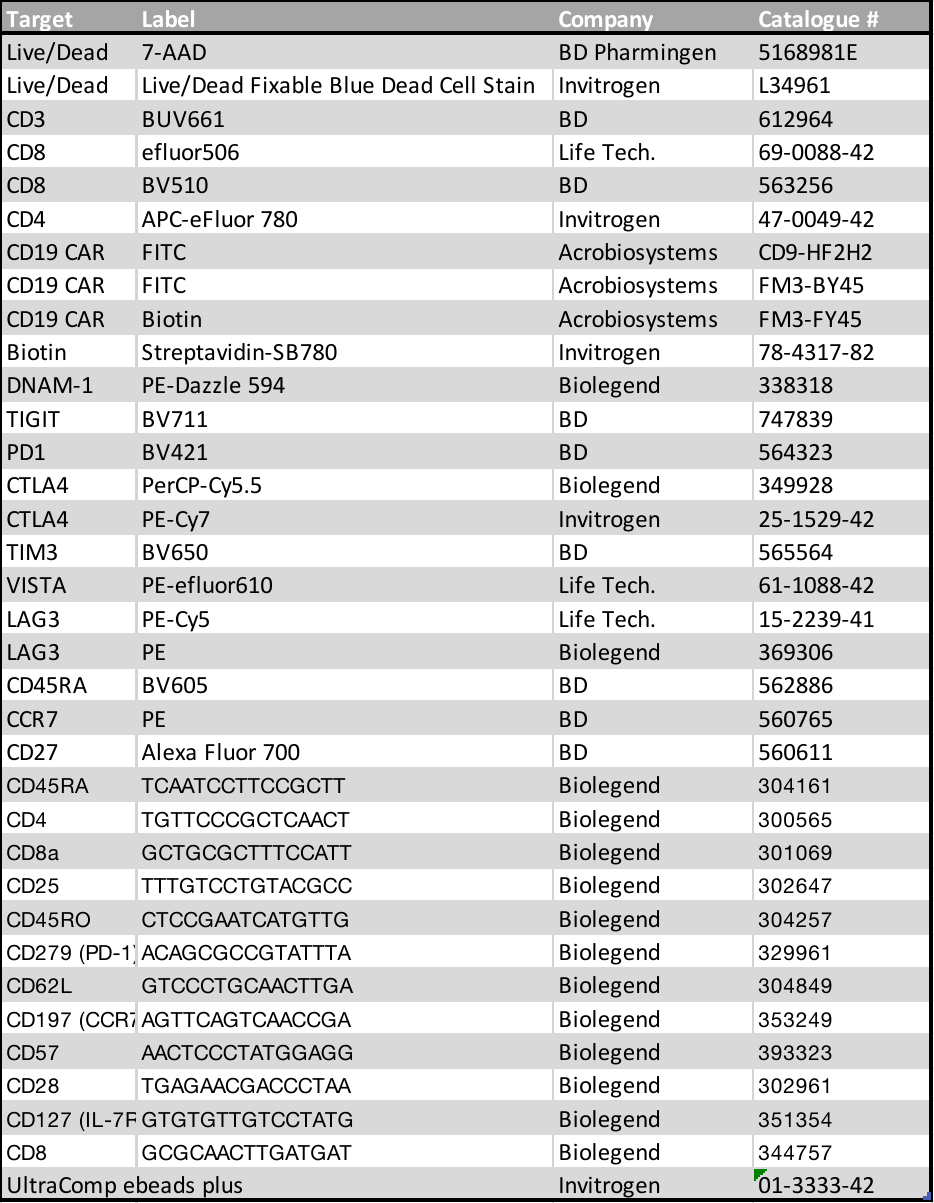

**Supplemental Figures**

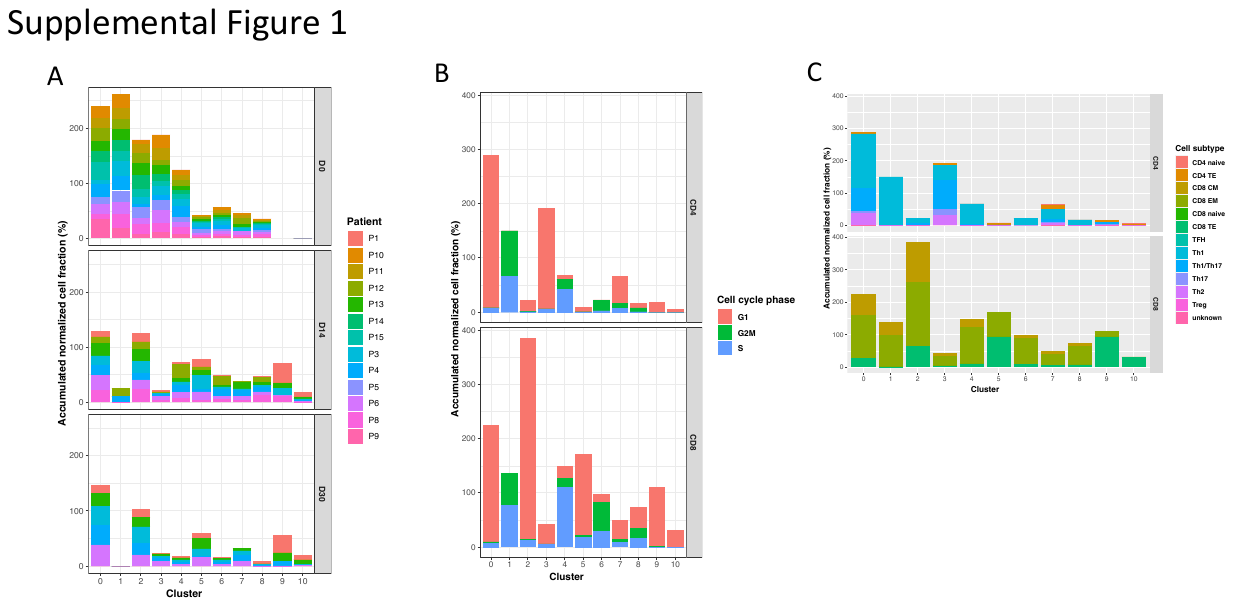

**Supplemental Figure 1. CD19 CAR-T cells demonstrate significant transcriptional heterogeneity that changes after infusion into patients. A)** Normalized proportions of CAR-T cells from each patient sample in each cluster separated by time point. Some patients are not represented at day 14 or day 30. **B)** Normalized proportions of CAR-T cells in the cell cycle phase of each cluster separated by CD4 and CD8 cell types. **C)** Normalized proportions of CD4 and CD8 CAR-T cell subtypes in each cluster separated by CD4 and CD8 cell types.

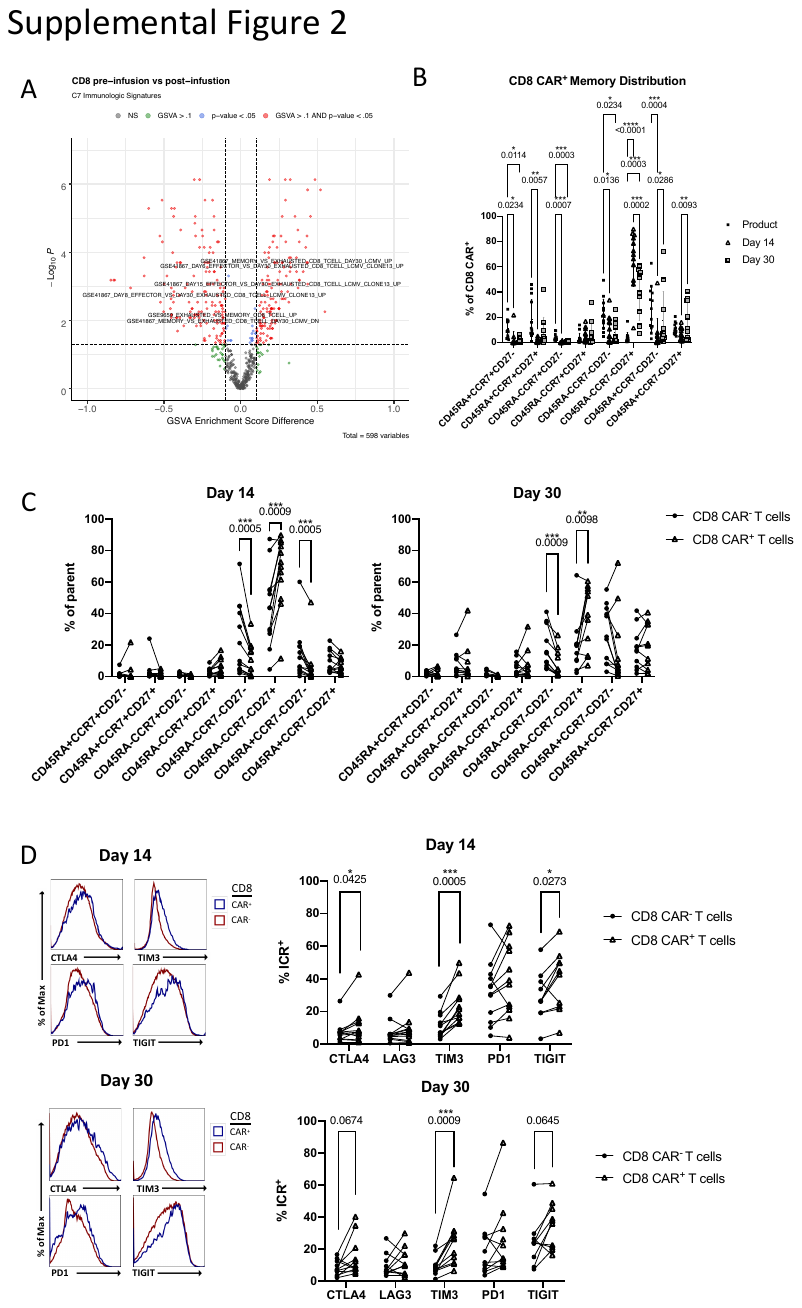

**Supplemental Figure 2. Circulating CD8 CAR-T cells differentiate to an effector-like state and express high levels of TIGIT post-infusion. A)** Volcano plot of gene set enrichment (p-value < 0.05) between post-infusion CD8 CAR-T cells compared to product CD8 CAR-T cells. Gene sets were taken from C7 signatures and filtered on “CD8”. **B)** Percentage of all memory marker combinations in CD8 CAR-T cells between the product, day 14, and day 30 time points as measured by flow cytometry. **C)** Percentage of all memory marker combinations in endogenous CD8 T cells compared to CD8 CAR-T cells at day 14 and day 30 as measured by flow cytometry. **D)** Left – Histograms of fluorescence intensity of exhaustion markers between endogenous CD8 T cells compared to CD8 CAR-T cells as measured by flow cytometry. Each curve represents a concatenation of all samples with equal proportions of CD8 T cells from each sample. Right – Comparison between endogenous CD8 T cells and CD8 CAR-T cells of the percentage of cells expressing checkpoint receptors CTLA4, LAG3, TIM3, PD1, or TIGIT as measured by flow cytometry.

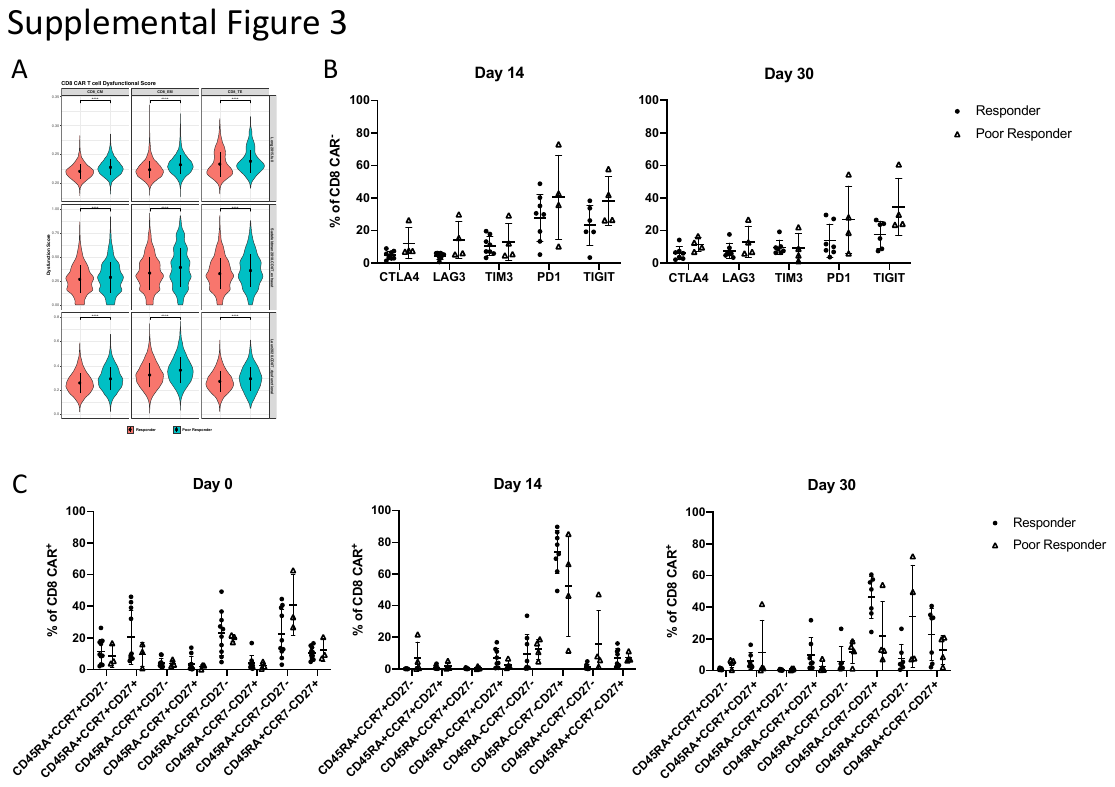

**Supplemental Figure 3. CAR-T cells of poor responders are enriched in an exhaustion-like phenotype post-infusion with high TIGIT expression. A)** Violin plot comparison dysfunction scores between response groups with three gene sets within effector memory, central memory, or terminal effector CD8 cell subtype assignments. **B)** Percentage of endogenous CD8 T cells expressing exhaustion markers CTLA4, LAG3, TIM3, PD1, or TIGIT between response groups as measured by flow cytometry. **C)** Comparison between response groups of memory markers co-expressed in CD8 CAR-T cells at each time point as measured by flow cytometry.

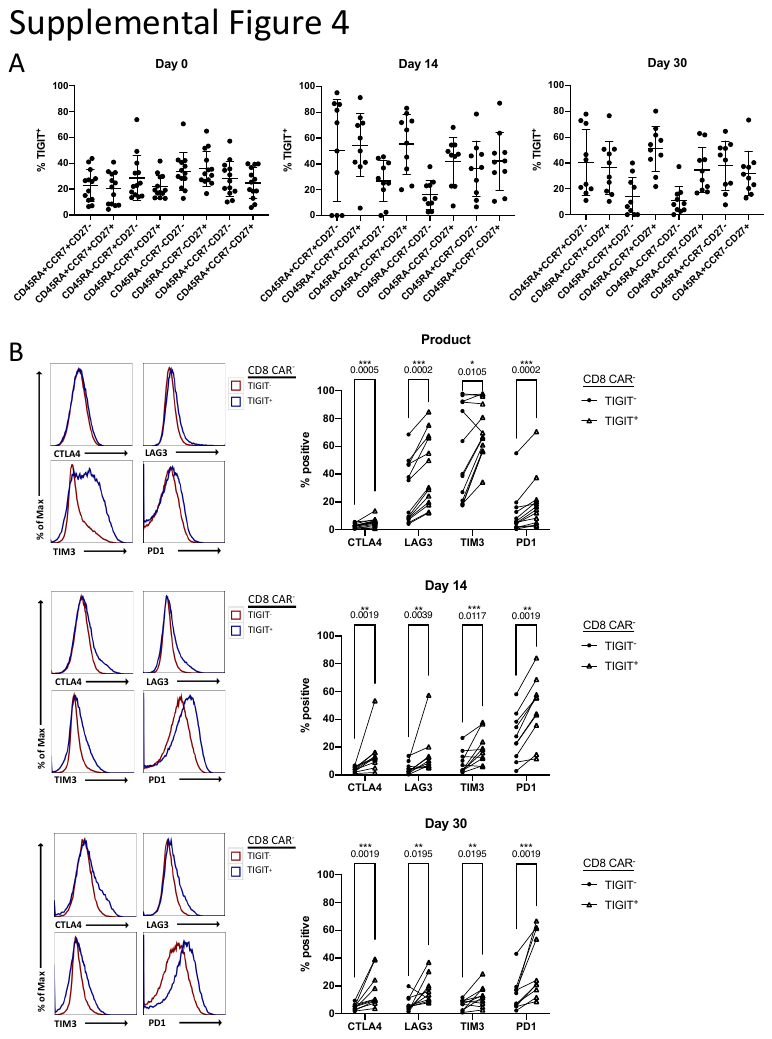

**Supplemental Figure 4. TIGIT expression is increased in CAR-T cells with an exhaustion phenotype. A)** Memory markers co-expressed in TIGIT^+^ CD8 CAR-T cells at each time point as measured by flow cytometry. **B)** Left – Histograms of fluorescence intensity of exhaustion markers between TIGIT^+^ and TIGIT^-^ CD8 CAR-T cells as measured by flow cytometry. Each curve represents a concatenation of all samples with equal proportions of CD8 T cells from each sample. Right – Comparison between TIGIT^+^ and TIGIT^-^ CD8 CAR-T cells of the percentage of cells expressing checkpoint receptors CTLA4, LAG3, TIM3, or PD1 as measured by flow cytometry.
